# Influence of Response Criterion on Nociceptive Detection Thresholds and Evoked Potentials

**DOI:** 10.1101/2021.11.02.466896

**Authors:** Boudewijn van den Berg, L. Vanwinsen, G. Pezzali, Jan R. Buitenweg

## Abstract

Pain scientists and clinicians search for objective measures of altered nociceptive processing to study and stratify chronic pain patients. Nociceptive processing can be studied by observing a combination of nociceptive detection thresholds and evoked potentials. However, it is unknown whether the nociceptive detection threshold measured using a Go-/No-Go (GN) procedure can be biased by a response criterion. In this study, we compared nociceptive detection thresholds, psychometric slopes and central evoked potentials obtained during a GN procedure with those obtained during a 2-interval forced choice (2IFC) procedure to determine 1) if the nociceptive detection threshold during a GN procedure is biased by a criterion and 2) to determine if nociceptive evoked potentials observed in response to stimuli around the detection threshold are biased by a criterion. We found that the detection threshold can be higher when assessed using a GN procedure in comparison with the 2IFC procedure. The average P2 component in the central evoked potential showed on-off behavior with respect to stimulus detection and increased proportionally with the detection probability during a GN procedure. These data suggest that nociceptive detection thresholds estimated using a GN procedure are subject to a response criterion.

## 1 Introduction

Pain scientists and clinicians search for objective criteria to identify impaired nociceptive processing for the purpose of stratification and treatment of chronic pain patients (Mouraux & Jannetti, 2018). With this aim, nociceptive processing of patients is usually evaluated using a combination of neurophysiological and psychophysical testing. In this field, there is a recent renewed interest in the assessment of mechanical, thermal and electric detection thresholds. However, the interpretation of these thresholds could alter depending on the procedure through which these thresholds are measured.

Recently, we developed a method to assess nociceptive processing by quantifying the effect nociceptive stimulus properties on detection probability and cortical evoked potentials (EPs). In this method, we stimulate nociceptive afferents in the skin by intra-epidermal electric stimulation with a specialized electrode (Steenbergen et al., 2012). This method selectively activates nociceptive afferents in the skin provided that low stimulation currents are used, for which a limit of twice the detection threshold was proposed as a rule of thumb (Mouraux, 2010). Stimulus amplitudes are centered around the detection threshold by an adaptive psychophysical method of limits (Doll, Veltink, & Buitenweg, 2015) and the electroencephalogram (EEG) is recorded in response to each stimulus. This allows us to record the combination of nociceptive detection thresholds and evoked potentials in response to nociceptive stimulation. We recently showed that nociceptive detection thresholds of single-pulse and double-pulse intra-epidermal electric stimuli can be used to observe peripheral and central changes of nociception following deafferentation by capsaicin (Doll et al., 2016). Nociceptive evoked potentials can be used as a marker for altered central nociception, e.g. in central sensitization (van den Broeke et al., 2015), attentional modulation (Legrain, Guérit, Bruyer, & Plaghki, 2002) or placebo analgesia (Wager, Matre, & Casey, 2006). The combination of both methods allowed us to evaluate the effect of temporal stimulus properties on nociceptive detection threshold and evoked potentials in healthy participants (van den Berg & Buitenweg, 2021; van den Berg et al., 2020), and could be used to study impaired nociceptive processing in chronic pain patients in future studies.

Although nociceptive detection thresholds appear sensitive to induced changes in peripheral and central nociceptive processing, it remains unclear how observed detection threshold are related to the underlying physiological systems. In all of our studies, we have used an adaptive method of limits with a Go-/No-Go (GN) procedure to approach and estimate the detection threshold, i.e.: 1) an adaptive series of stimuli is presented, 2) the participant has to indicate when a stimulus was detected and 3) the stimulus amplitude is increased or decreased depending on stimulus detection. Subsequently, logistic regression was used to estimate the detection threshold and slope based on all available data. Although the obtained detection threshold is used to probe central or peripheral nervous function, most studies appear to disregard the fact that these thresholds could also be modulated by a sensory, perceptual or decision criterion (Georgeson, 2012). In addition, it still remains unknown how this response criterion is related to evoked brain activity, measured in some studies as a more ‘objective’ measure of altered nociceptive or somatosensory processing. In this work, our aim was to determine how this criterion dependency affects the results obtained during measurements of nociceptive detection thresholds and evoked potentials.

The role a response criterion can be omitted by using a 2-interval forced choice (2-IFC) procedure (Kingdom & Prins, 2016), where participants are asked to choose during which of two observation intervals a stimulus was applied. During the 2IFC procedure, the interval is reported correct if the sample during the interval with a stimulus is larger than the sample during the interval with only noise. In this study, we compared nociceptive detection thresholds, psychometric slopes and central evoked potentials obtained during a GN procedure with those obtained during a 2IFC procedure with two objectives. Our first objective was to determine if the nociceptive detection threshold during a GN procedure is biased by a response criterion, i.e. resulting in a different detection threshold with respect to the 2IFC threshold. Our second objective was to determine if a bias of the detection threshold by a response criterion is reflected in the nociceptive evoked potentials observed in response to stimuli around the detection threshold.

## 2 Methods

The results presented in this work include measurements of the detection threshold using a GN and a 2IFC procedure in randomized order. A total of 25 participants was included and performed both procedures. In the last 15 participants, also the EEG was recorded during task performance. The experiments were performed at the University of Twente, the Netherlands, and were approved by the local Medical Review and Ethics Committee. All experiments were performed in accordance with the declaration of Helsinki.

### 2.1 Participants

A total of 25 healthy participants (12 males and 13 females, age 19-30) were included in this study. The inclusion criterion was an age between 18 and 40 years old. Exclusion criteria were skin abnormalities at the site of stimulation, diabetes, implanted stimulation devices, pregnancy, usage of analgesics within 24 hours before the experiment, the consumption of alcohol or drugs within 24 hours before the experiments, pain complaints at the time of the experiment, a medical history of chronic pain or any language problems that would impede communication with the participant. All participants provided written informed consent before participation in the experiment.

### 2.2 Stimuli

Each stimulus consisted of cathodic square wave electric pulses generated by a constant current stimulator (NociTRACK AmbuStim, University of Twente, Enschede, The Netherlands). Stimuli were delivered to the epidermis at the back of the right hand via a custom made electrode consisting of 5 inter-connected microneedles protruding 0.5 mm from the electrode surface. Intra-epidermal electric stimulation preferentially activates nociceptive afferents in the skin, provided that stimuli remain below twice the detection threshold (Mouraux, 2010; Poulsen, Tigerholm, Meijs, Andersen, & Mørch, 2020). A previous validation study of the electrode used in this study demonstrated that electric pulses resulted in a sharp pricking sensation (Steenbergen, 2012). Two stimulus types were used during the experiment:

- One square pulse with a pulsewidth of 210 μs.
- Two square pulses with a pulsewidth of 210 μs and an inter-pulse interval of 10 ms.

### 2.3 Familiarization

Participants were instructed to press and hold a button. For familiarization with the sensation of intra-epidermal stimuli, participants were stimulated with a series of pulses with a stepwise (0.025 mA) increasing amplitude and instructed to release the button when a stimulus was clearly perceived for at least two times. For an initial estimate of the detection threshold for each stimulus type, participants were stimulated with a series of pulses with a stepwise (0.025 mA) increasing amplitude and instructed to release the button when any sensation was perceived that they ascribed to stimulation.

### 2.4 Go/No-Go Procedure

Participants were seated upright in a chair and asked to focus on the site of stimulation. Detection thresholds were estimated and tracked using an adaptive procedure (Doll et al., 2015). Participants were instructed to press and hold a button, and to briefly release the button when any sensation was perceived that they ascribed to stimulation (Fig. 1). For the adaptive procedure, the stimulus amplitude was randomly picked from a vector of 5 stimulus amplitudes with a stepsize of 0.025 mA initialized around the initial estimate of the detection threshold. The vector of amplitudes was decreased by 0.025 mA when a stimulus was reported as detected and increased by 0.025 when the participant did not release the response button. This process was repeated independently for every stimulus type, with the order of stimulus type randomized, for a total of 130 stimuli per type.

**Figure 1:**
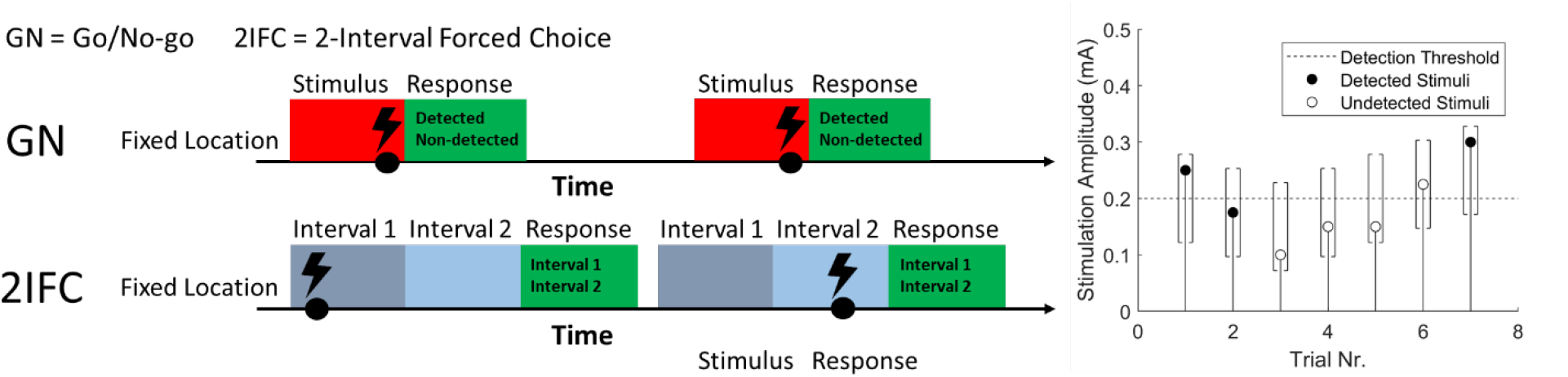
The Go/No-Go and the 2-Interval Forced Choice (2IFC) procedure used measure the nociceptive detection threshold for single-and double pulse intra-epidermal electric stimuli (left). The adaptive procedure used to converge to the detection threshold with a vector of 5 stimulus amplitudes from which the stimulus is randomly selected. When a stimulus is not detected, the vector of 5 stimulus amplitudes (with a stepsize of0.025 mA) is decreased by 0.025 mA (Go-/No-Go) or 0.075 mA (2IFC). When a stimulus is detected, the vector of 5 stimulus amplitudes is increased by 0.025 mA (both procedures).

### 2.5 2-Interval Forced Choice Procedure

Participants were seated upright in a chair and asked to focus on the site of stimulation. Detection thresholds were estimated and tracked using an adapted version of the adaptive procedure in previous version. Participants were stimulated during one of two time intervals (Fig. 1), marked by an auditory cue. After each set of two time intervals, participants were asked to indicate during which time interval they were stimulated. For the adaptive procedure, the stimulus amplitude was randomly picked from a vector of 5 stimulus amplitudes with a stepsize of 0.025 mA initialized around the initial estimate of the detection threshold. The vector of amplitudes was decreased by 0.075 mA when the reported time interval was incorrect and increased by 0.025 when the reported time interval was correct. Note that the decrease after an incorrect answer [*d_incorrect_*] is 3 times larger than the increase after a correct answer [*d_correct_*], as this ratio is governed by the value of the detection threshold, 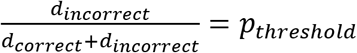, where *p_threshold_* is equal to 0.5 for a GN procedure and equal to 0.75 for a 2-IFC procedure. This process was repeated independently for every stimulus type, with the order of stimulus type randomized, for a total of 130 stimuli per type.

### 2.6 Electroencephalography

The scalp EEG was recorded at 32 channels (international 10/20 system) using a REFA amplifier (TMSi B.V, Oldenzaal, the Netherlands) with a sampling rate of 1024 Hz. Participants were asked to fix their gaze at a spot on the wall. Electrode impedance was kept below 20 kΩ.

### 2.7 Nociceptive Detection Threshold

The nociceptive detection probability was estimated by global optimization of the negative log­ likelihood using an implementation of the GlobalSearch algorithm (Ugray et al., 2007) in combination with an interior-point algorithm to find local minima (Coleman & Li, 1996) in Matlab. In the case of the Go-/No-Go procedure (Equation (1)), the detection probability was modeled using a cumulative normal distribution as a function of an intercept [*β*_0_], additive temporal summation of the first pulse [*β*_*A*1_ *A*] and the second pulse [*β*_*A*2_ *A*], and a linear drift over time [*β_t_t*]. In the case of a 2IFC procedure (Equation (2)), this function was adapted to account for a 50% guessing rate at low stimulus amplitudes.

Detection probability for a go/no-go procedure:

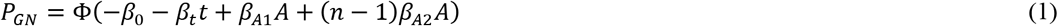

Detection probability for a 2-interval forced choice procedure:

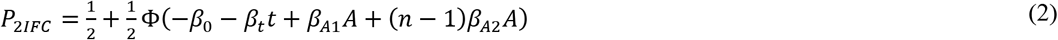

### 2.8 Evoked Brain Activity

The EEG was preprocessed using the FieldTrip toolbox (Oostenveld, Fries, Maris, & Schoffelen, 2011). Eye-blinks, eye movement and movement artefacts were corrected using independent component analysis (Delorme, Sejnowski, & Makeig, 2007). Epochs with excessive EMG activity or remaining movement artefacts were removed by visual inspection. Grand average EP waveforms of detected and non-detected stimuli (GN), and of correct and incorrect stimuli (2-IFC) were computed at T7-F4 and CPz-M1M2 and tested for significance with respect to baseline and with respect to the other condition using cluster-based nonparametric permutation tests (Maris & Oostenveld, 2007). In addition, grand average EP waveforms were computed for four levels of detection probability (.00-.25, .25-.50, .50-.75 and .75-1.0) for both procedures. Significance of the effect of detection probability on the EEG was assessed by fitting a linear mixed model (3) to the EEG at each latency, and obtaining the t-value of effect coefficients using Satterthwaite’s approximation of the degrees of freedom. The t-values were corrected for retesting over time using the Benjamini-Hochberg correction (Hochberg & Benjamini, 1995). The average P2 amplitude for each of the four levels of detection probability was determined by averaging over time between 380 ms and 420 ms post-stimulus.

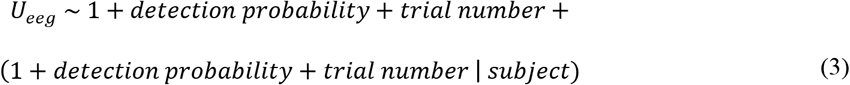

## 3 Results

### 3.1 Nociceptive Detection Threshold

A typical example of an experiment with the GN and the 2-IFC procedure is displayed in Fig. 2. During the GN procedure, the detection threshold for single-pulse stimuli was larger than the detection threshold for double-pulse stimuli. Both thresholds showed a small increasing drift over time. During the 2-IFC procedure, the thresholds were equal for single-pulse and double-pulse stimuli. Drift over time was small or not present. The detection thresholds and slopes for all 25 participants are displayed in Fig. 3. The detection threshold for single-pulse stimuli during a 2-IFC procedure was significantly lower than the detection threshold for single-pulse stimuli during a GN procedure. The psychometric slope for single-pulse stimuli during a 2-IFC procedure was significantly larger than the psychometric slope for single-pulse stimuli during a GN procedure.

**Figure 2:**
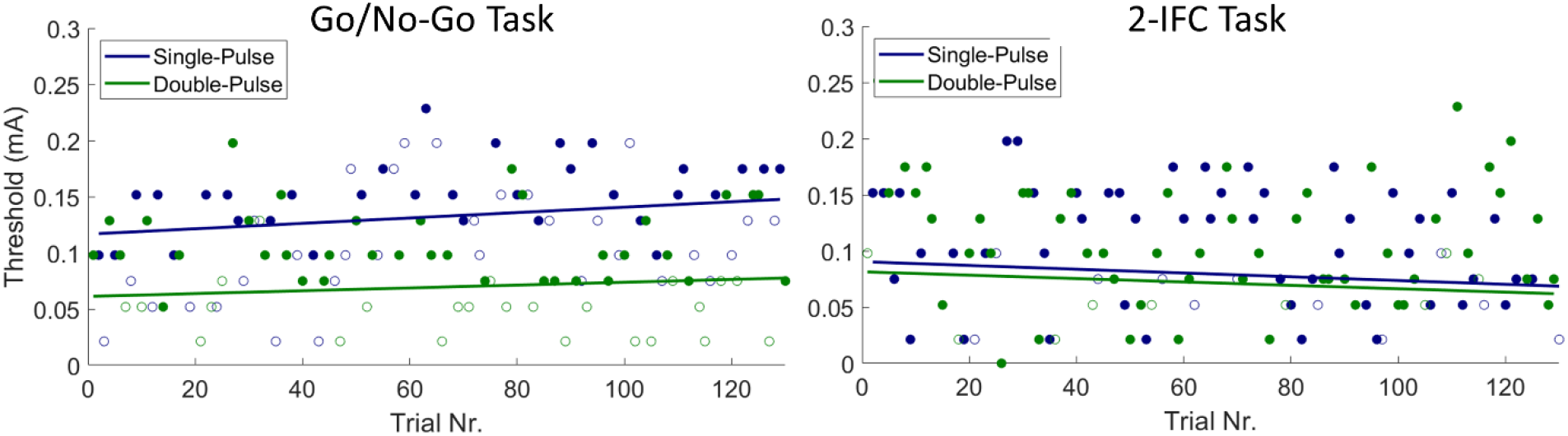
Typical example of detection thresholds obtained when peiforming a GN procedure (left) and when peiforming a 2IFC procedure (right). When peiforming a 2-IFC procedure, detection thresholds appeared to equalize for both stimulus types. Detected and non-detected (GN) or correct and incorrect (2IFC) stimuli are depicted by closed and open circles respectively.

**Figure 3:**
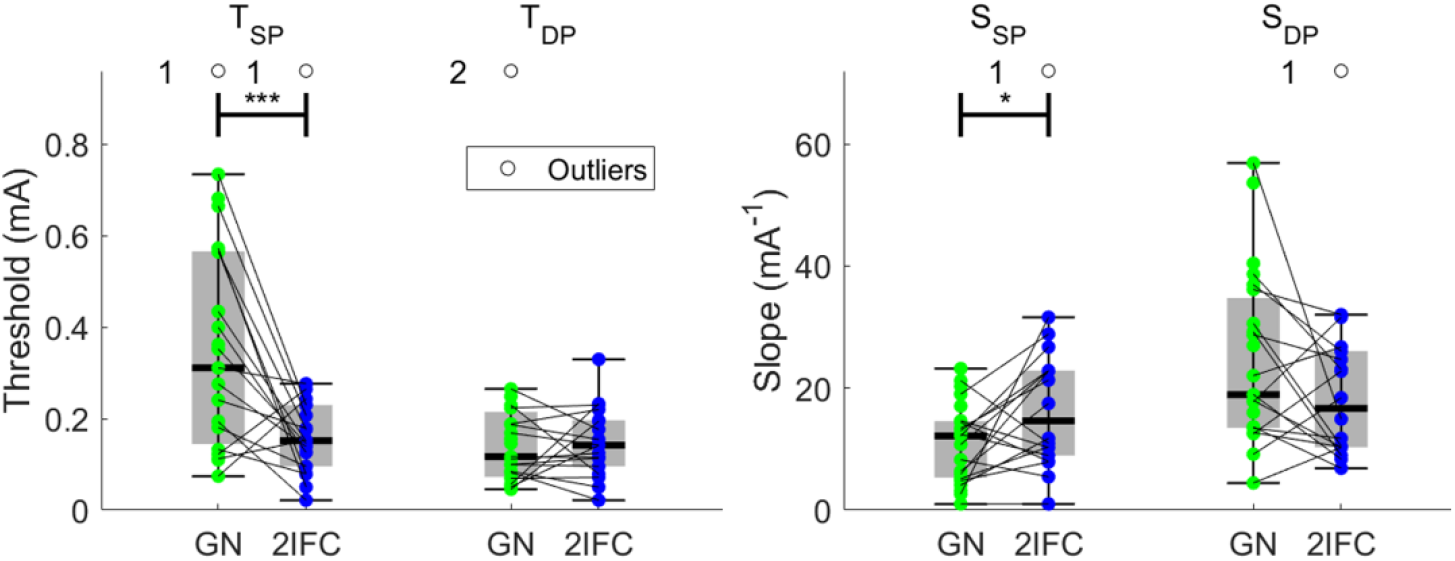
Individual results and boxplots of the detection thresholds (T) for single-pulse (SP) and double-pulse (DP) stimuli during the go-/no-go (GN) and the 2-interval forced choice (2IFC) procedure for all 25 participants. Significance is indicated with * (p<.05), ** (p<.01) and *** (p<.001). Detection thresholds for single-pulse stimuli were significantly lower and psychometric slopes were significantly larger when assessed in a 2IFC procedure in comparison with the GN procedure.

### 3.2 Evoked Brain Activity

Grand average evoked potentials at Cz-M1M2 acquired during both procedures are displayed in Fig. 4. There was a significant contrast between evoked potentials in response to detected and non­ detected stimuli in the GN procedure, and correct and incorrect trials in the 2IFC procedure. For the GN procedure, the evoked potential was significantly larger than baseline for detected as well as non-detected stimuli. For the 2IFC procedure, the evoked potential was only significantly larger than baseline for correct trials. Note that the average evoked potential for correct trials (2IFC) was lower than the average evoked potential for detected stimuli (GN), but might be confounded by inclusion of trials that were not consciously perceived but simply guessed correctly.

**Figure 4:**
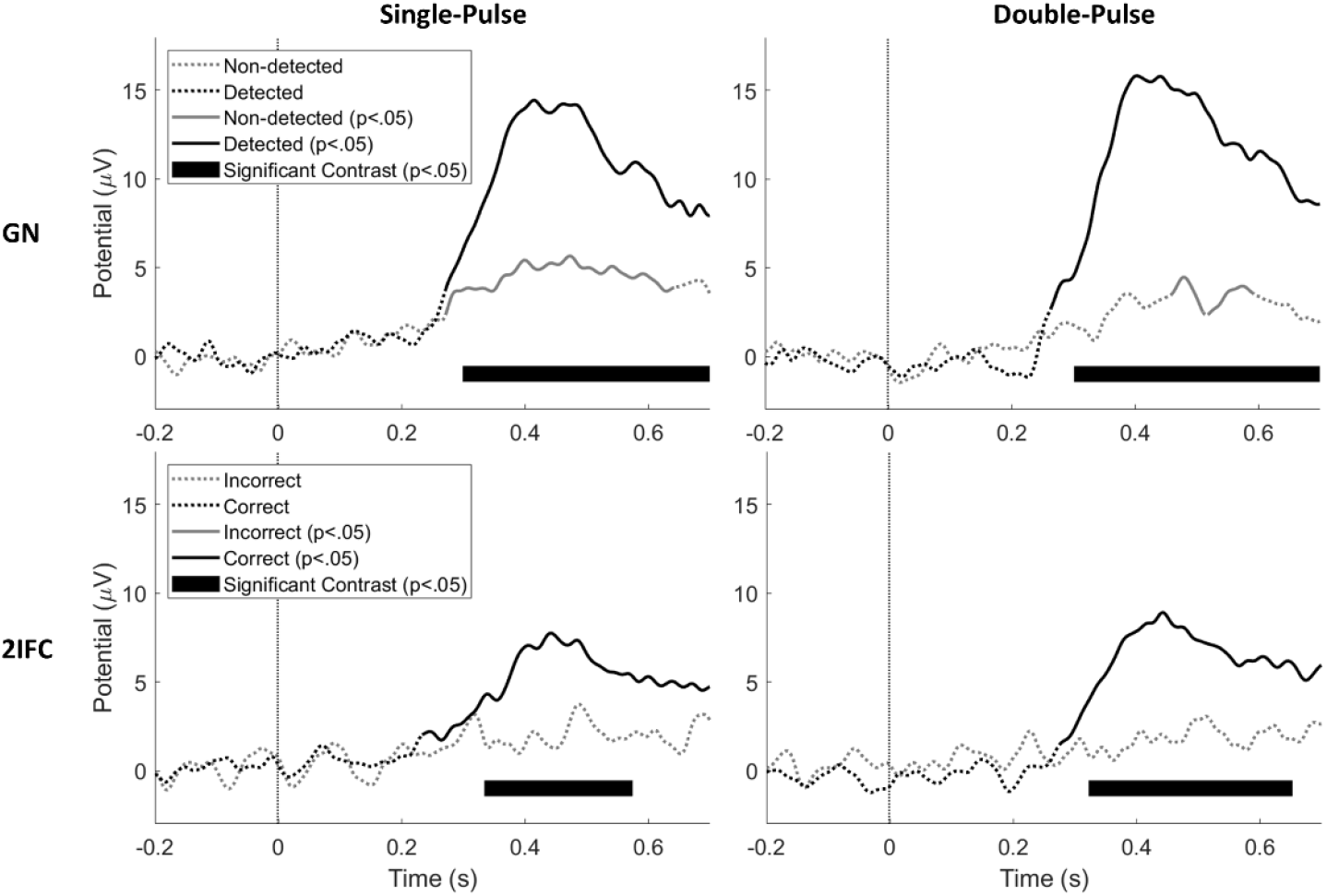
Grand average evoked potential at Cz-MJM2 for stimuli acquired during both procedures. Significance with respect to baseline (p<.05) is indicated by a solid line, while insignificant parts are indicated by dotted lines. Latencies with a significant contrast (p<.05) between detected and non-detected (in GN procedure) and between correct and incorrect (in 2IFC procedure) are marked with a black bar.

Grand average evoked potentials at Cz-MIM2 for several levels of detection probability are displayed in Fig. 5. There was a significant effect of detection probability on the evoked potential during both procedures and for both stimulus types. While the average evoked potential during a GN procedure appears graded with stimulus intensity, the average evoked potential during a 2IFC procedure remains low until high levels of detection probability are reached, i.e. a detection probability larger than 0.875. Both phenomena are more clearly visible in Fig. 6, where the average amplitude of the major positive peak between 380 and 420 ms, the P2, is displayed. Here, the average P2 appears to increase almost proportional with respect to detection probability during the GN procedure. Note that this proportional increase with detection probability can be attributed to two phenomena: 1) The average P2 for detected stimuli is at almost every point significantly larger than the average P2 amplitude for non-detected stimuli, leading to an increased average P2 over all stimuli when more stimuli are detected. 2) There is an increasing trend in the average P2 for both detected and non-detected stimuli, leading to a further increase in the average P2 over all stimuli with respect to detection probability. Similar to previous figure, the average P2 during the 2IFC procedure remains low until a probability larger than 0.875 is reached.

**Figure 5:**
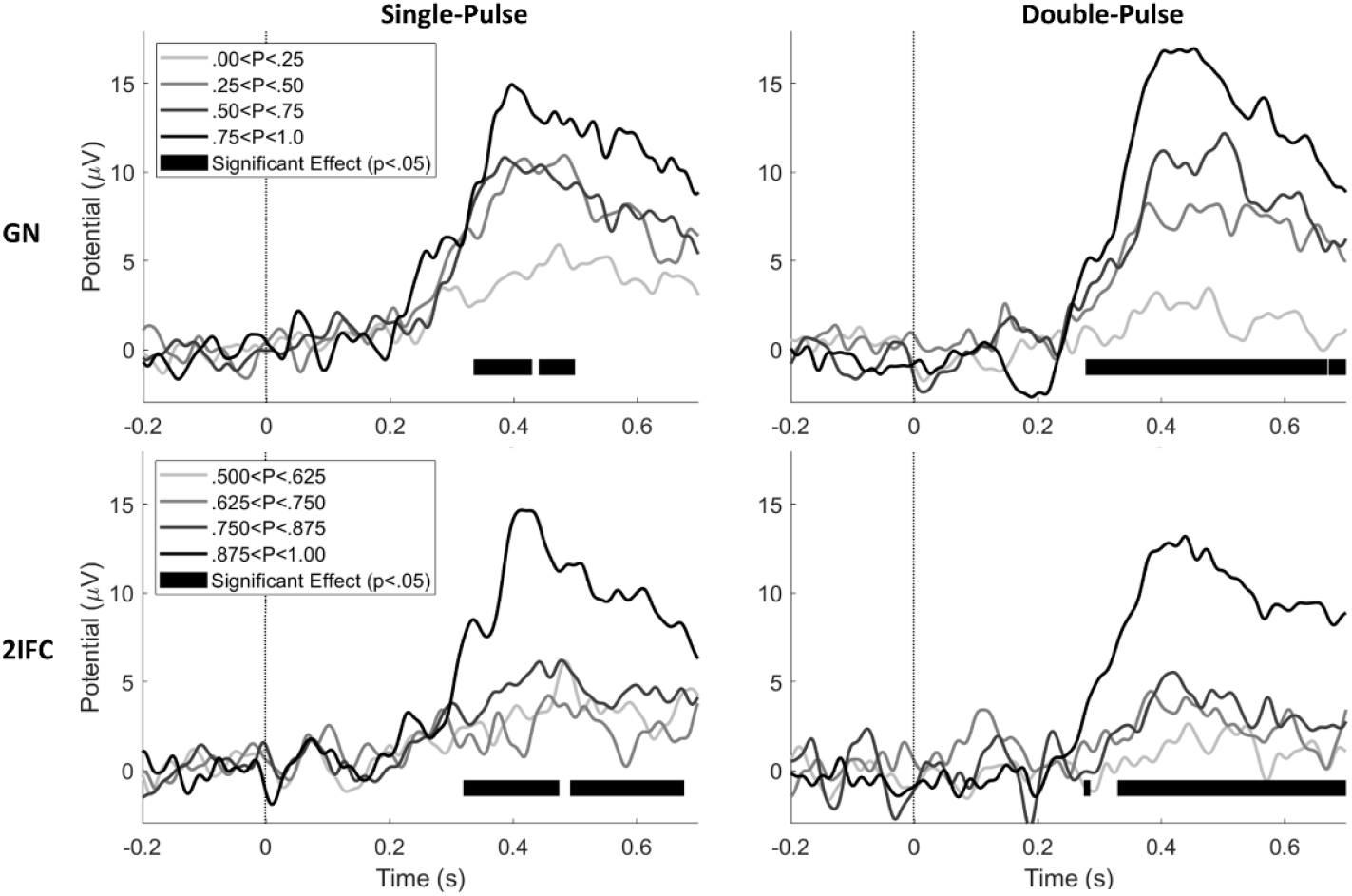
Grand average evoked potential at Cz-MJM2 for stimuli acquired during both procedures at 4 levels of detection probability. Latencies with a significant effect of detection probability (p<.05) are marked with a black bar.

**Figure 6:**
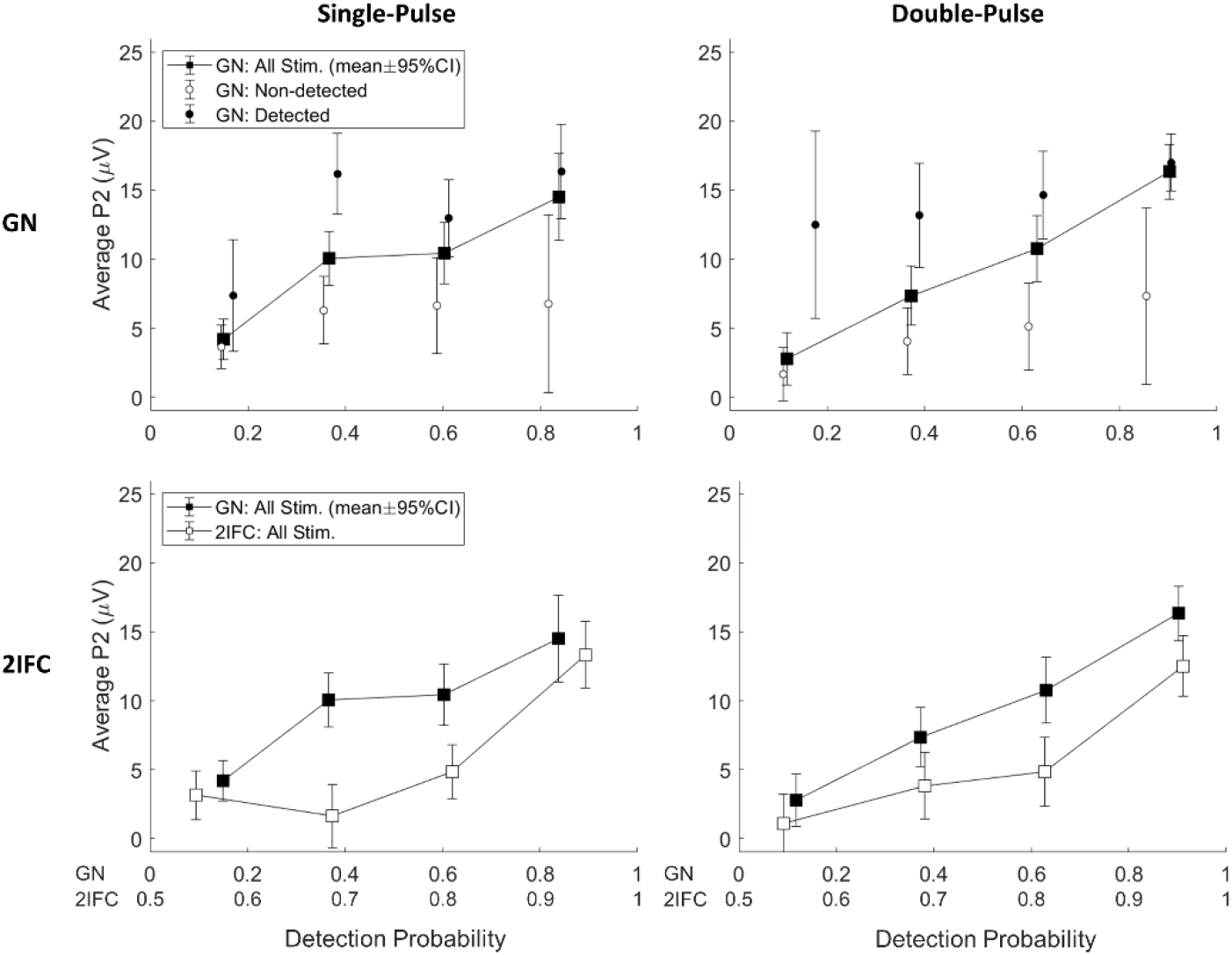
Average amplitude of P2 peak in the evoked potential (and 95%CI) with respect to detection probability. There is an almost proportional relation between the average P2 amplitude and detection probability. Average P2 amplitude is significantly larger for detected stimuli in comparison with non-detected stimuli, and shows an increasing trend with respect to detection probability for both detected and non-detected stimuli. In the 2IFC procedure, the average P2 amplitude remains at very low levels (comparable with or even lower than non-detected stimuli) until a high detection probability (>0.8 for GN and >0.9 for 2IFC) is reached.

## 4 Discussion

In this study, we observed nociceptive detection thresholds, psychometric slopes and central evoked potentials obtained during a GN and a 2IFC detection procedure. The differences observed between both procedures in nociceptive detection threshold and in evoked responses include important clues about how nociceptive detection might work, and how the threshold obtained during these procedures can be interpreted.

The first objective of this study was to determine if the nociceptive detection threshold during a GN procedure is biased by a response criterion. We found that the detection threshold for single-pulse intra-epidermal electric stimuli is significantly higher, and the psychometric slope significantly lower, during a GN procedure in comparison with a 2IFC procedure. In contrast, we found that the threshold for double-pulse stimuli does not differ significantly between procedures. This result implies that for some types of stimuli the nociceptive detection threshold measured during a GN procedure reflects evoked neural activity exceeding a response criterion, rather than the presence of sensory evidence itself Equal detection thresholds for double pulse stimuli between the GN and the 2IFC procedure indicate that the extend to which the observed detection threshold is influenced by the response criterion also depends on stimulus properties, and that the bias of the detection threshold introduced by a criterion might be lower for high signal-to-noise ratio stimuli such as the double-pulse stimulus in this experiment. In addition, a significant difference was observed between single- and double-pulse stimuli during a GN procedure, while no significant difference was observed between detection thresholds for single- and double-pulse stimuli during a 2IFC procedure. Although a small difference between the single- and double-pulse threshold might go unnoticed due to estimation errors, it is clear that the large difference between both stimulus types in a GN procedure almost completely disappears during 2IFC. The reason for this discrepancy between both tasks remains unclear without more sophisticated psychophysical modeling, which is out of the scope of this study. However, these results warrant the development of novel psychophysical models that are tailored to the process of nociception in future studies. One of the potential factors that might help explaining such a difference would be the presence of spontaneous neural activity influencing both the response criterion and psychometric slope of the participant. More importantly, formulation of psychophysical models that are connected to neurophysiological mechanisms can lead to more insight in the interpretation of the detection thresholds measured in a clinical or research setting.

The second objective of this study was to determine if the presence of a response criterion is reflected in the nociceptive evoked potentials observed in response to stimuli around the detection threshold. We measured a significant central evoked response at Cz-M1M2 during both procedures for detected stimuli (GN) and correctly reported trials (2IFC). We also measured a significant evoked response to non-detected stimuli (GN), which was absent for incorrectly reported trials (2IFC). We found that the evoked P2 response is proportionally graded with detection probability during a GN procedure. At the same time, we observed that the P2 response during a 2IFC procedure for stimuli with the same detection probability (corrected for guessing rate), remains low until a large detection probability is reached. The P2 response to detected and non-detected stimuli show that we might be looking at a mostly dichotomous response, where the response is much larger for detected stimuli than for non-detected stimuli. The visual evoked P3 response is considered a key marker of conscious access to sensory evidence (Rutiku, Martin, Bachmann, & Aru, 2015; Salti, Bar-Haim, & Lamy, 2012), and the high degree of overlap in activated brain regions suggests a similar functional significance of the nociceptive P2 (Jannetti & Mouraux, 2010; Mouraux & Jannetti, 2009). Our observation that the P2 shows an on-off behavior with respect to reported conscious perception is in accordance with this theory.

Assuming that we are looking at an entirely dichotomous response, we can explore how the detection probability in both procedures relates to the probability of evoking a central brain response at Cz-M1M2. Fig. 7 shows that the difference between detection probability and evoked response probability determines the observed pattern of the average P2 response in Fig. 6. When there is no difference between the detection threshold and the threshold for evoking a brain response at 0.5 probability, both curves will overlap leading to a proportional relation between the evoked response probability (or average P2) and the detection probability, as we observed for the GN procedure. When the detection threshold is lower than the threshold for evoking a brain response, we expect a bended curve which predicts that the evoked response probability (or average P2) remains low until a high detection probability is reached, as we observed for the 2IFC procedure. As such, our results suggest that the evoked response probability is equal to the detection probability in the GN procedure, but lower than the detection probability in the 2IFC procedure, implying that the central P2 brain response assessed in this experiment was evoked only when the stimulus exceeded the response criterion. When considering the nociceptive P2 as a marker for conscious access to sensory evidence, the response criterion observed in this experiment could be interpreted as a perceptual criterion, i.e. only stimuli above this criterion are perceived. This also implies that the average P2 responses observed during a GN procedure are affected by a response criterion just like the participant responses itself, when they are not corrected for stimulus detection.

**Figure 7:**
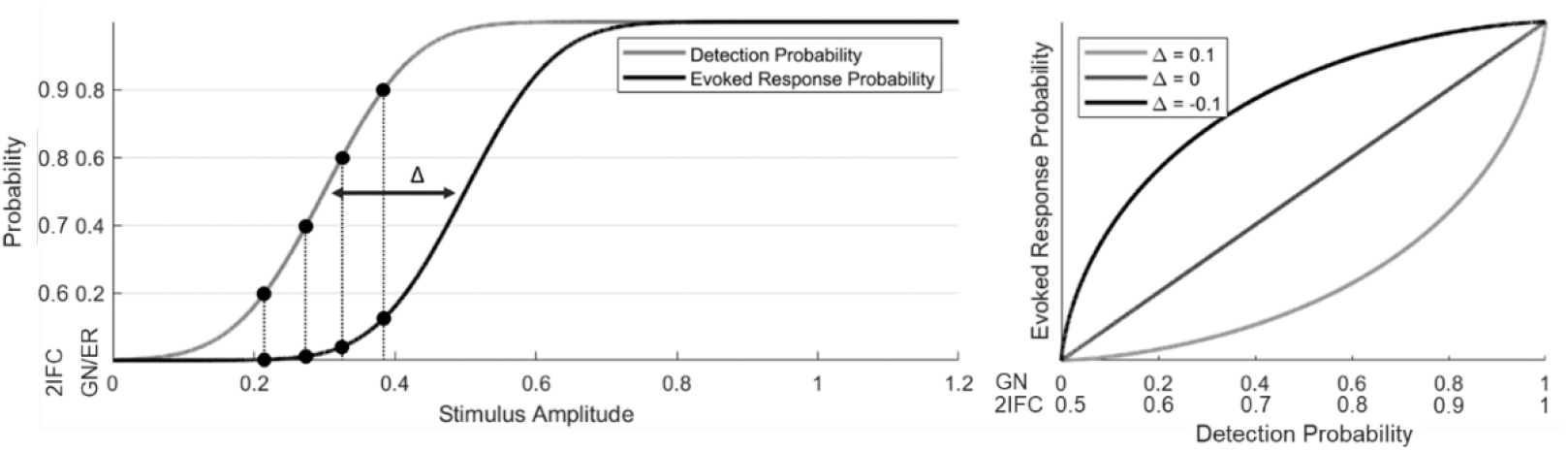
The difference between detection probability and evoked response probability determines the relation between detection probability and average evoked response.

These observations show that when the nociceptive detection threshold is assessed using a GN procedure, one might observe the effect of an adjusted perceptual criterion rather than altered nociceptive processing following an intervention. This has important consequences for studies using nociceptive detection thresholds to assess altered central and peripheral nociceptive processing. The mechanical and thermal detection threshold are increased in patients with neuropathic pain and signs of central sensitization (Maier et al., 2010). Thermal heat and cold detection thresholds show a high sensitivity to potential peripheral nerve damage by diabetes (Courtin et al., 2020) as well as painfulness in diabetic neuropathies (Krämer, Rolke, Bickel, & Birklein, 2004). Intra-epidermal electric detection thresholds are increased following deafferentiation by capsaicin (Doll et al., 2016) and following diabetic neuropathy (Suzuki et al., 2016). Our current results emphasize that these nociceptive detection thresholds can in some cases reflect a central criterion that determines if the stimulus is consciously perceived, rather than the threshold for activation of the nociceptive system itself This criterion does not only affect participant report, but also the central P2 response, which appeared to be generated only when the stimulus was reported as consciously perceived, i.e. when the stimulus exceeded a perceptual criterion. The notion that we can measure the potential influence of a perceptual criterion by comparing detection thresholds in a GN and 2IFC procedures opens up new avenues of research into the role of perception in nociceptive processing and (chronic) pain.

## 5 Declarations

### 5.3 Ethics Approval

All experiments were approved by the local ethics committee and in accordance with the declaration of Helsinki.

### 5.4 Consent to Participate

All participants provided written informed consent and were rewarded for participation in the experiment.

### 5.5 Consent for Publication

All authors approved the manuscript and agree with submission of this preprint to bioRxiv.

### 5.6 Availability of Data and Materials

A limited dataset of the experiments reported here is available on request.

### 5.7 Code Availability

Code required to perform the analyses reported here is available on request.

